# Insights on the effect of mega-carcass abundance on the population dynamics of a facultative scavenger predator and its prey

**DOI:** 10.1101/2023.11.08.566247

**Authors:** Mellina Sidous, Sarah Cubaynes, Olivier Gimenez, Nolwenn Drouet-Hoguet, Stéphane Dray, Loïc Bollache, Daphine Madhlamoto, Nobesuthu Adelaide Ngwenya, Hervé Fritz, Marion Valeix

**Affiliations:** Chrono-environnement UMR6249, CNRS, Université Bourgogne Franche-Comté, F-25000, Besançon, France; CEFE, Univ Montpellier, CNRS, EPHE, IRD, Montpellier, France; CNRS, Université de Lyon, Université de Lyon 1, VetAgroSup, LBBE, UMR 5558, F-69622, Villeurbanne, France; Long-Term Socio-Ecological Research Site (LTSER) France, Zone Atelier “Hwange”, Hwange National Park, Bag 62, Dete, Zimbabwe; OFB, 5 All. de Bethléem, 38610 Gières, France; Zimbabwe Parks and Wildlife Management Authority, Main Camp Research, Hwange National Park, Zimbabwe; REHABS International Research Laboratory, CNRS – Université Lyon – Nelson Mandela University, George Campus, Madiba Drive, 6531 George, South Africa; Sustainability Research Unit, Nelson Mandela University, George Campus, Madiba Drive, 6531 George, South Africa

## Abstract

The interplay between facultative scavenging and predation has gained interest in the last decade. The prevalence of scavenging induced by the availability of large carcasses may modify predator density or behaviour, potentially affecting prey. In contrast to behavioural mechanisms through which scavenging affects predation, the demographic effects of facultative scavenging on predator and prey populations remain poorly studied. We used the semi-natural experimental opportunity in Hwange National Park, Zimbabwe, where contrasted management measures (culling and artificial supply of water) have led to fluctuations in elephant carrion abundance, to identify the consequences of facultative scavenging on the population dynamics of a large mammalian carnivore, the spotted hyaena (*Crocuta crocuta*), and its prey. Using a 50-year dataset and Multivariate Autoregressive State Space models, we estimated hyaena and prey densities over four periods contrasted in elephant carrion availability due to management practices. Models that allow hyaena and their prey populations’ growth rate to vary depending on these four periods contributed significantly to explain variations in their density, which is consistent with an effect of management measures on the population dynamics of hyaena and its prey. Although our results support a predominant role of bottom-up mechanisms, whereby hyaena density is driven by herbivore density, itself driven by resources availability, some subtle patterns of densities could be interpreted as consequences of changes in predation pressure following changes in scavenging opportunities. We discuss why signals of prey and predator population dynamics decoupling are less likely to be observed in systems with a high diversity of prey, such as African savannas, and why inputs of mega-carcasses as pulsed resources hardly impacted top-down relationships in the long run. This study represents a first investigation of the long-term effects of carrion pulses, whose frequency may increase with climate changes, on the classical predator-prey coupling for large mammals.

## Introduction

Predation is the most studied ecological process amongst those linking predator and prey populations (Abrams 2000, Guiden et al. 2019). Although overlooked, scavenging (i.e., when animals feed on dead animals and carrion they did not kill) recently gained increased recognition (Wilson and Wolkovich 2011, Moleón and Sánchez-Zapata 2015, Luna et al. 2021). So far, these two processes have been little investigated together. However, because carnivores are often facultative scavengers (Fallows et al. 2013, Pereira et al. 2014, Gomo et al. 2017), and scavenging is energetically less costly than hunting, carnivores are distracted from their live prey in the presence of carrion (DeVault et al. 2003). Consequently, carrion and live prey are linked by indirect interactions. On the one hand, live prey can be subject to a higher predation pressure, or hyper-predation, if carrion pulses lead to a higher abundance of carnivores (through demography or local immigration) but carrions are not available (because of interference or decay of the carrion; Courchamp et al. 2000, Cortés-Avizanda et al. 2009a, Cortés-Avizanda et al. 2009b, Moleón et al. 2014). On the other hand, live prey can be subject to a reduced predation pressure, or hypo-predation, if carnivore abundance remains stable but the rate of consumption of live prey is reduced and compensated by scavenging (Bate and Hilker 2012, Fallows et al. 2013, Moleón et al. 2014, Mellard et al. 2021). These indirect interactions linking carrion and live prey, through predators, have been studied theoretically (Andrén et al. 2011, Mellard et al. 2021), experimentally (Cortés-Avizanda et al. 2009a), and empirically (Cortés-Avizanda et al. 2009b, Fallows et al. 2013). These previous works globally conclude about an impact of scavenging on predation processes, either through an alteration of prey and predator space use (Cortés-Avizanda et al. 2009b), or through a direct modification of predation rate (Cortés-Avizanda et al. 2009a, Andrén et al. 2011, Fallows et al. 2013, Mellard et al. 2021). However, there is a dearth of long-term, demographic studies to assess whether regular pulses of carrion interfere with the population dynamics of predators and their prey (Moleón et al. 2014). Still, in some systems, population dynamics of a meso-predator was affected by facultative scavenging with associated consequences on prey populations (Roth 2003), and regular anthropogenic food subsidies seem able to alter the classical predator-prey coupling of population dynamics (Rodewald et al. 2011). The megafauna is unparalleled in the extent of carrion they provide after death and mega-carcasses (from whales, elephants, rhinoceros) can be considered as important pulsed resources for scavengers.

In this study, we explored the effect of elephant (*Loxodonta Africana*) carrion availability on the numerical response of a large opportunist scavenging predator, the spotted hyaena (*Crocuta crocuta*; hyaena hereafter; Figure 1; Kruuk 1972) and its prey. Their relative abundances were monitored over almost 50 years in Hwange National Park, Zimbabwe. As a consequence of management practices, this ecosystem witnessed large variations in the abundance of the elephant population over the years, and four periods characterized by different elephant carcass availability emerged.

**Figure 1.**
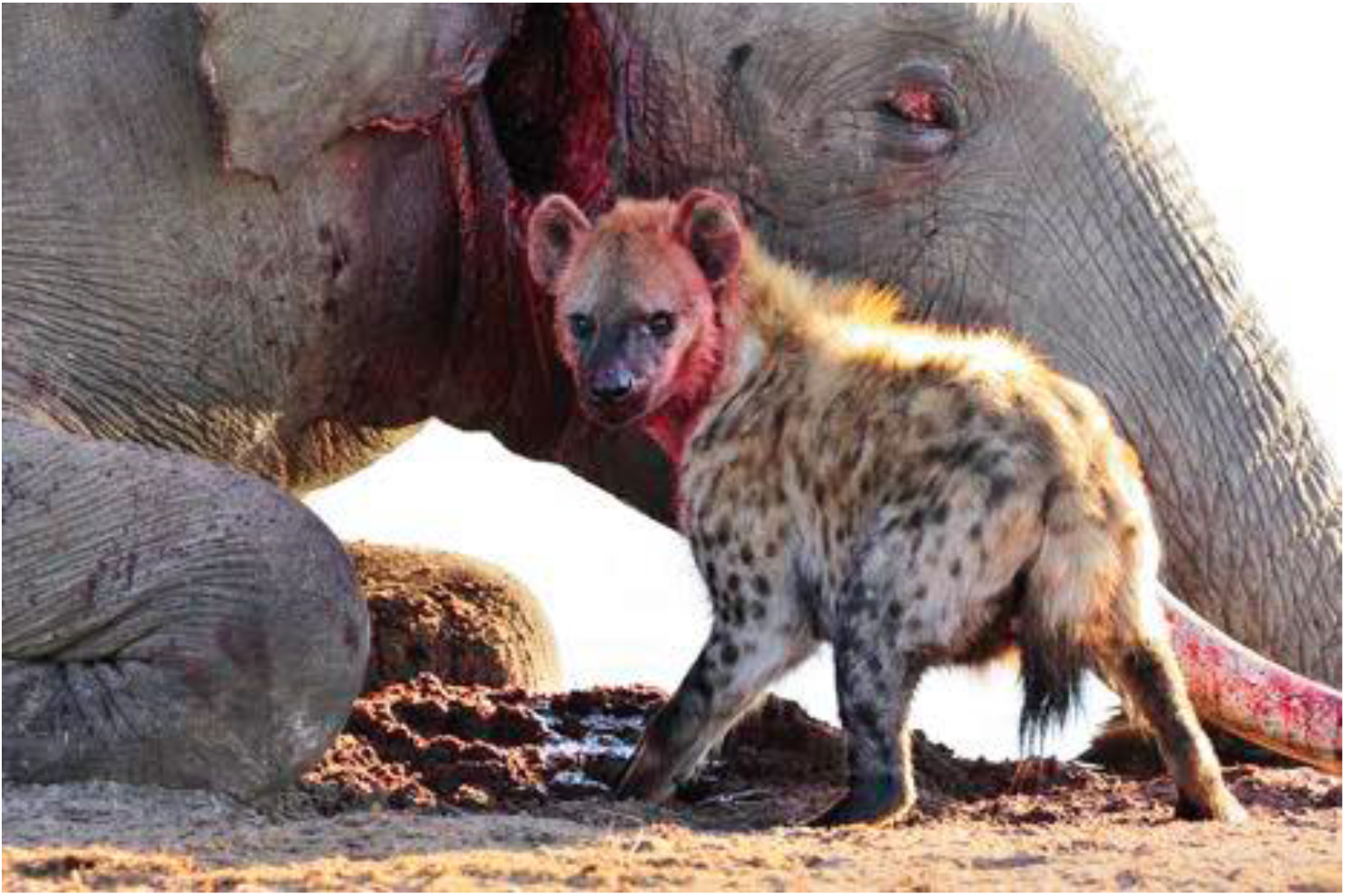
Hyaena scavenging from an elephant carcass (credit: S. Périquet).

From 1960 to 1986, massive culls were undertaken in the area in response to an elephant abundance perceived as “too high” regarding its effects on biodiversity (Child 2004, Slotow et al. 2008). During this time, at least 18,000 elephants were killed (Cumming 1981, ZPWMA 1998, Child 2004, Slotow et al. 2008). This period corresponds to our first study period (1972-1986) when hundreds to a few thousand carcasses were available annually and clearly provided an important source of food for scavengers. In contrast, in the second period (1987-1992), elephant carcasses were rare due to a steady and continuous increase of the population triggered by surface water management (pumping of underground water) and the cessation of culling (Chamaillé-Jammes et al. 2008). This lasted until the third period (1993-2008) when elephant carcasses were present in large numbers in dry years (Dudley et al. 2001), after the elephant population reached density-dependence, fluctuating around 30,000 individuals (Chamaillé-Jammes et al. 2008). Finally, during the fourth period (2009-2020), natural mortality (density-dependence driven) still existed but was likely to be much lower than during period 3 due to the creation of additional waterholes (Appendix 1), allowing the elephant population to pass the equilibrium reached previously and start growing again.

The existence of these four periods contrasted in terms of elephant carcass availability, together with the long-term monitoring of hyaena and prey populations covering these different periods, provide a unique semi-natural experimental opportunity to assess the impacts of scavenging on predation at the level of population dynamics. Here, we hypothesised that if scavenging modified predation with further demographic consequences, we would detect that the Hwange hyaena population is driven by its live prey in periods of low elephant carcass abundance (periods 2 and 4), but less dependent of its live prey in periods of high elephant carcass abundance (periods 1 and 3).

## Methods

### Study area

Hwange National Park covers ~15,000 km^2^ of dystrophic semi-arid (~600 mm annual rainfall) savanna in western Zimbabwe (19°00’ S, 26°30’ E). This protected area is not fenced, which allows wildlife to move freely. Surface water is primarily found in natural depressions that hold rainwater. However, most dry up as the dry season progresses, from May to September. To guarantee sufficient drinking water for animals in the dry season, water has been managed since 1935 through the pumping of underground water into waterholes distributed mostly in the northern sector of the park. The number of pumped waterholes has varied over the past 50 years, from about 20 to more than 80 (Appendix 1). In this study, we focus on a 3,000 km^2^ study area in the north-eastern sector of the park (Figure 2), the most touristic area of the park, with most pumped waterholes.

**Figure 2.**
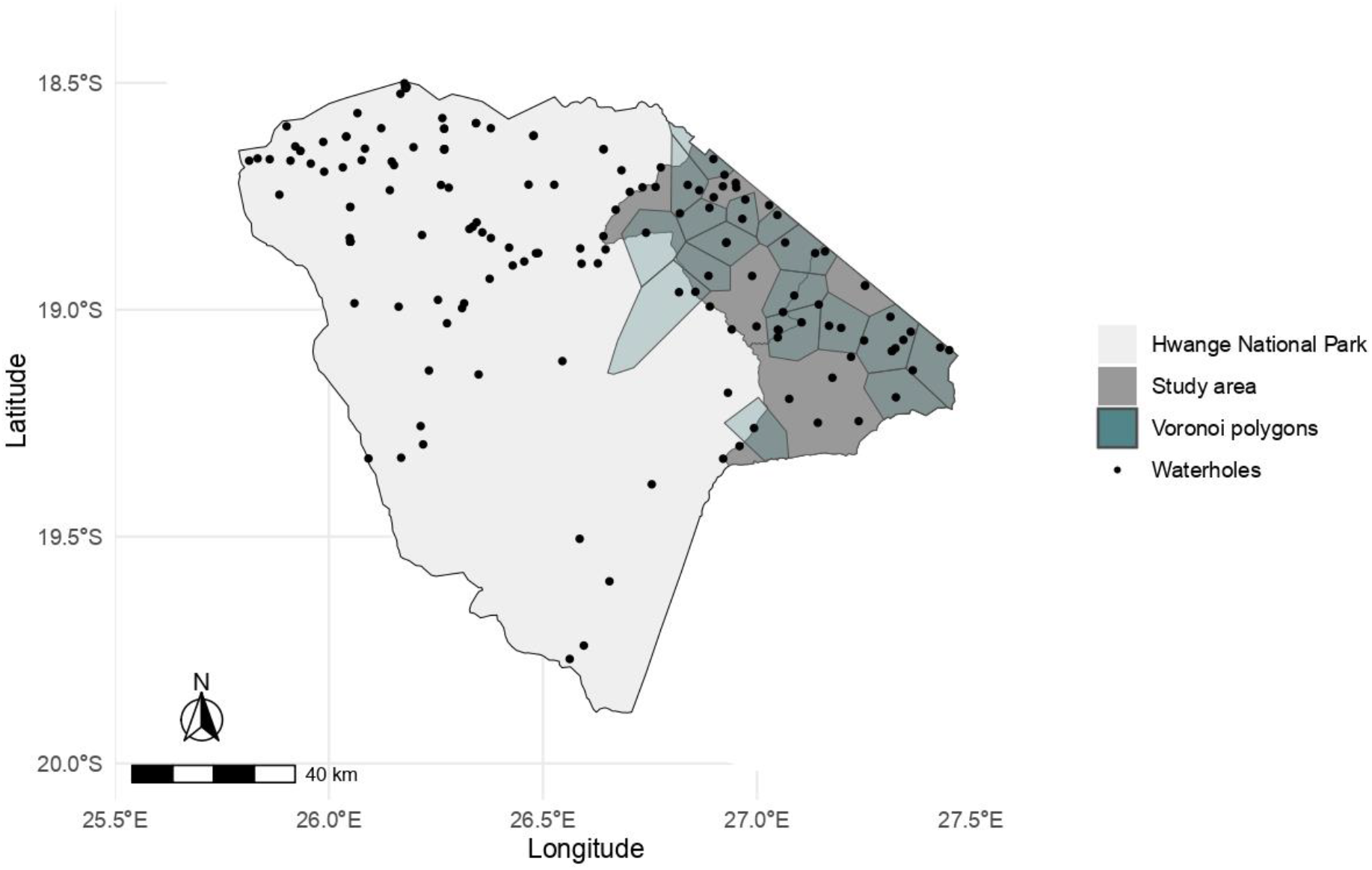
Hwange National Park and location of the study area. Black dots are the waterholes of the park. Voronoi polygons used for the analysis are coloured in blue.

### Data

We used a long-term data set resulting from the regular monitoring of waterholes from 1972 to 2020 by the Wildlife Environment Zimbabwe (WEZ). This monitoring consists of annual 24h counts during full moon at the end of the dry season. At this time of the year, it is mostly the pumped waterholes that provide drinking water to animals, but in years with a good rainy season a few natural pans can hold water too. Because this monitoring is largely influenced by annual rainfall, counts are characterized by high inter-annual variance and cannot be considered as total counts. However, abundance trends detected with these counts are the same as those obtained with other more traditional methods (aerial census; Valeix et al. 2008), and thus we consider that long-term trends inferred from counts at waterholes reliably reflect actual patterns of abundance changes in populations (as in Valeix et al. 2008). Additionally, to control for the proximity of some waterholes and the differential surfaces considered as their attraction zones, monitored waterholes were grouped in new spatial units created using a centroidal Voronoi tessellation based on pumped waterholes (Okabe et al. 2009) (Figure 2). Indeed, if we consider a pumped waterhole, its attraction is reduced when new pumped waterholes are added in its vicinity or in years when neighbouring natural pans still hold water. In other words, we expect less animals to be observed at a focused pumped waterhole since animals should split between this waterhole and the neighbouring pumped or natural waterholes. This could lead to conclude that species abundance has decreased when in reality the patterns observed come from this change in attraction rather than an actual decrease in regional abundance of species. By grouping neighbouring waterholes into new spatial units, we expect to reduce this bias arising from the fluctuation in the number of waterholes over years. For each year, we summed the animals counted at the different waterholes of each spatial unit and divided it by the number of waterholes surveyed, and then by the surface of the spatial unit to assess densities. For political (civil war) and economic reasons, counts were not performed in 1974, 1983, and from 1976 to 1981. For the analyses, we selected the spatial units that were monitored for at least 50% of the years composing each period, except for period 1 (the period impacted by the civil war) for which we kept units with at least 25% of the years monitored.

One spatial unit was excluded for the analyses because it is known for the presence of a hyaena den close to a waterhole and thus presented singularly high hyaena counts. Counts were log-transformed and we dealt with 0 values by adding 0.0001 to the whole dataset, which corresponds to one order of magnitude lower than the smallest value of the data.

We used data on hyaena and its prey. For prey, we pooled all (males, females, adults, sub-adults and juveniles) preferred and common prey species of hyaena in Hwange National Park (Périquet et al. 2015) (‘prey’ hereafter). This includes plains zebra (*Equus quagga*), greater kudu (*Tragelaphus strepsiceros*), sable (*Hippotragus niger)*, roan *(Hippotragus equinus*), blue wildebeest (*Connochaetes taurinus)*, waterbuck (*Kobus ellipsiprymnus*), impala (*Aepyceros melampus*), and warthog (*Phacochoerus africanus*).

### Statistical analyses

We fitted a Multivariate Autoregressive State-Space Model (Holmes et al. 2012) using R package MARSS (Holmes et al. 2021, R Core Team 2021) to estimate prey and hyaena densities from 1972 to 2020 from counts data at the waterholes, while accounting for observation and process errors. We considered each spatial unit to be a replicate of a common dynamic, and therefore estimated one density for prey and one density for hyaena for the entire study area. MARSS separates the state process, capturing variations in species density, from the observation process explaining variations in the number of individuals observed at the waterholes. More precisely, the model uses an autoregressive process, which means that the state x of the system at time t (x_t_) depends on its value at time t-1 (x_t-1_). In our case, x is the densities of a guild (either prey or hyaena) and we have:

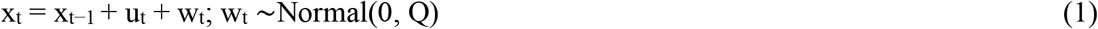

Where x_t_, x_t-1,_ and Q are 1×1 matrices. x_t_ and x_t−1_ represent the abundance estimated at time t and t-1 respectively for the focus guild. The parameter w_t_ is the error term of the state process and represents environmental stochasticity. This environmental stochasticity is modelled as a normal distribution with mean 0 and a variance Q that takes the same values for the whole time series of each guild. The parameter u_t_ is the growth rate of the population at time t. This is the parameter that was constrained differently in the three models we tested. For each time step, the growth rate has a unique value and thus u_t_ is a 1×1 matrix.

Equation (1) models the state process. Here, it represents the changes in densities of each guild over time. We have only access to a noisy signal of these dynamics through their observation at the different waterholes of our study area. The densities of each guild can thus be considered as “hidden states” and state-space models aim at retrieving access to these states through their signals. This is done by incorporating a second equation that models the observation process:

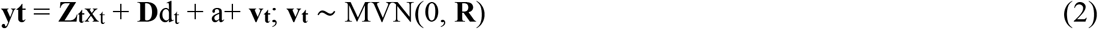

For n waterholes, **y**_**t**_ has a dimension nx1, where n is the number of waterholes. **Z** is a nx1 matrix that links the observations **y**_**t**_ to the state to be estimated (x_t_). Since we only estimated the density of each species separately and not in the same model, Z matrix is full of 1s. The bias parameter, a, is set to be equal between sites. The parameter v_t_ is the error term of the observation process equation, and comes from a multivariate normal distribution that share a common variance over time and site. The variance is the parameter **R** a variance covariance matrix of dimension *nxn*. Finally, d_t_ is a vector with annual rainfall value at year t, and **D** further links the effect of d_t_ to the **y**_**t**_ that are inputs of the model. In this observation model, we included annual rainfall as a covariate to account for bias on counts at waterholes (Appendix 2). During dry years, animals need to visit waterholes more often than during wet years when more ephemeral natural ponds can be found in the park. Thus, when annual rainfall is not incorporated into the model, estimated densities are overestimated in dry years, and underestimated in wet year (Appendix 2).

We fitted and compared (using AICc; Burnham et al. 2011) three different models: (i) a *constant* model with a constant growth rate, (ii) a *time-varying* model estimating a growth rate per year, and (iii) a *period* model estimating one growth rate per period. The period model aims at testing a signal in densities caused by the fluctuations of elephant carrion availability. We used an analysis of deviance (ANODEV) to test the significance and assess the proportion of the deviance explained by a period-dependent growth rate for hyaena and its prey (Grosbois et al. 2008).

## Results

Observed counts data and estimated population densities for prey and hyaena are provided in Figure 3. The period model best described the variations in density of prey (AICc=1862.05, 1890.89 and 1850.95 for the constant, time-varying and period models respectively) and hyaena (AICc=3039.18, 3089.02 and 3029.70 for the constant, time-varying and period models respectively). ANODEV tests were significant for both prey (F = 0.009; r^2^ = 0.22; df= 3) and hyaena (F= 0.002; r^2^ = 0.28; df = 3). Hyaena and prey densities showed similar trends across the 4 study periods: the two populations increased during period 1, stabilized during period 2, decreased during period 3, and increased again during period 4 (Figure 3; period models’ growth rate estimates in Table 1). Differences between growth rate of hyaena and prey are similar in both periods when elephant carcasses were less abundant (0.069 in the second and 0.063 in the fourth period; Table 1). During periods 1 and 3 characterized by increased abundance of elephant carrion in the landscape, this difference was increased or decreased (0.123 during the first and 0.004 during the third period). However, confidence intervals of the two guilds partially overlap and are large.

**Table 1:** Estimation of growth rate (with 95% confidence interval) from the period model for prey and hyaena populations.

|  | Prey | Hyaena |
| --- | --- | --- |
| **Period 1**<br>(1972-1986) | 0.052 (0.017; 0.092) | 0.175 (0.079; 0.269) |
| **Period 2**<br>(1987-1992) | -0.010 (-0.073; 0.049) | 0.059 (-0.094; 0.204) |
| **Period 3**<br>(1993-2008) | -0.042 (-0.063; -0.021) | -0.058 (-0.107; -0.008) |
| **Period 4**<br>(2009-2020) | 0.025 (-0.009; 0.056) | 0.090 (0.013; 0.168) |

**Figure 3.**
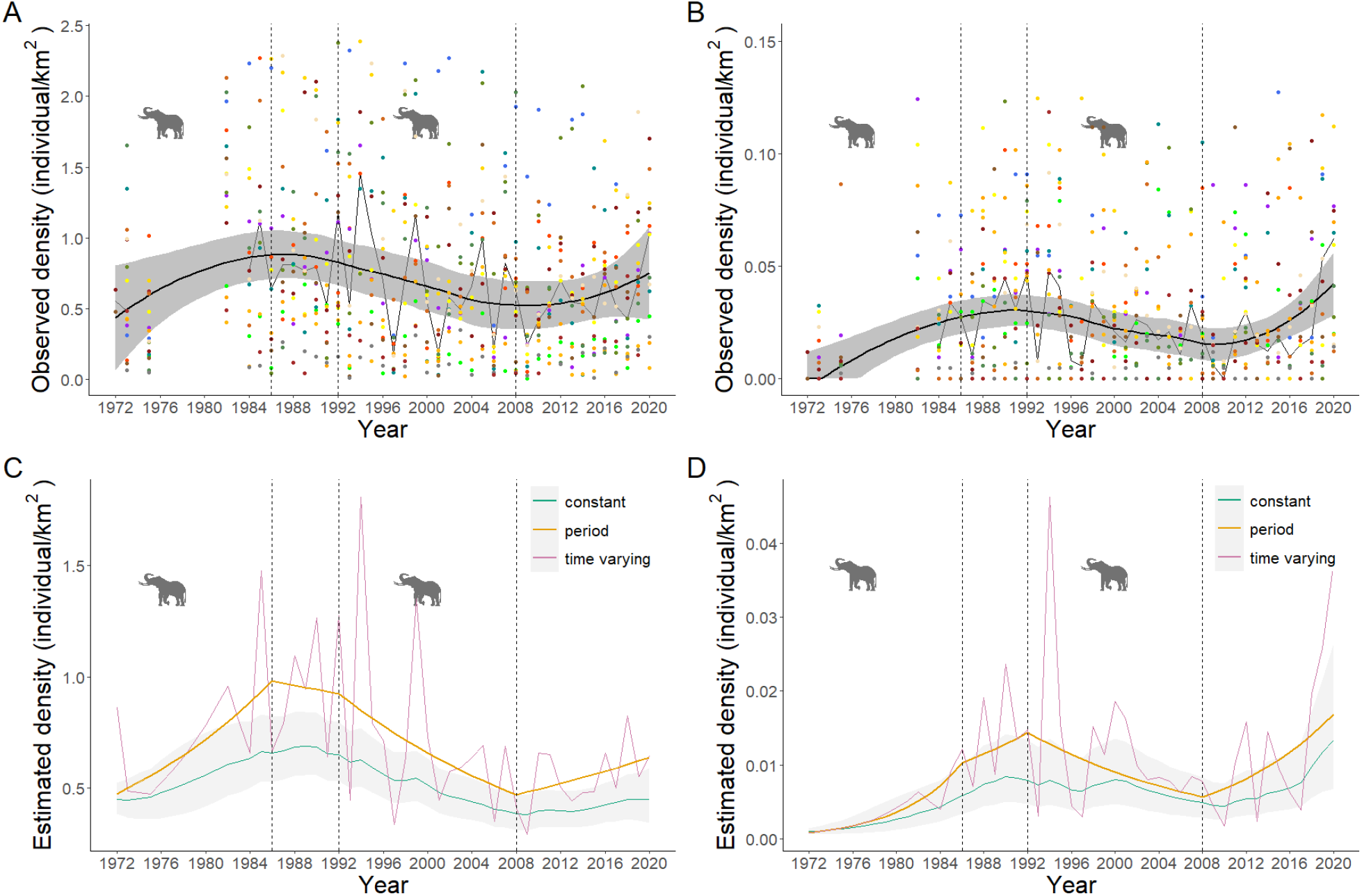
Top: Observed population densities of prey (**A**) and hyaena (**B**). Dots correspond to the densities observed at the spatial units (Voronoi polygons), each colour representing a spatial unit (95% of the data kept for visual clarity). Fluctuating thin lines represent the median of the observed densities each year. Bold line and ribbon represent the smooth of the median and its 95% confidence interval. **Bottom:** Estimated population densities of prey (**C**) and hyaena (**D**). Model estimates are given for the constant, time-varying, and period models. The ribbon represents the 95% of these estimates, but is very close to the estimates for the time-varying and period models. In all panels, the dotted vertical lines delimitate the four study periods; elephant pictogram indicates periods of high elephant carcass availability.

## Discussion

A megafauna’s carcass, which represents several tons of resource, undoubtedly influences the foraging and ranging behaviour of facultative scavengers, which can use such a carrion for long periods. In marine ecosystems, sharks can use whale carrion for several days (Fallows et al. 2013). In African savannas, hyaenas definitely modify their foraging and ranging habits in the presence of an elephant carcass, which they can use for up to two weeks (Cozzi et al. 2015, Périquet et al. 2021). During this time, hyaenas are diverted from their prey. In periods of high elephant carcass availability, these carrion-induced behavioural changes should affect interactions between carnivores and their prey. Surprisingly, little is known about the cumulative long-term effects of these changes and the ultimate implication for population dynamics of predators able of facultative scavenging and their prey. Using almost 50 years of data, and a semi-natural experiment involving carrion of the largest terrestrial mammal, we do not detect patterns supporting effects of high carcass availability on the population dynamics of hyaena and its prey (i.e. we do not detect that hyaena population is less driven by its live prey in periods of high elephant carcass abundance, as we initially hypothesized). The main result is that hyaena and its prey showed the same population trends over all periods, with hyaena population dynamics following that of its prey independently of elephant carcass availability, as in classical predator-prey fluctuations (Table 1 and Figure 3; Krebs et al. 2001, Hebblewhite et al. 2002, Wang et al. 2009).

The periods contributed significantly to explain variations in prey and hyaena density (~ 30% of the deviance). Thus, management measures (elephant culling and water supply) and the resulting elephant population dynamics used to define the periods partly explained population trends. Prey abundance increased in the first and last periods, when artificial waterholes (determining the amount of resource available for herbivores) were added in the landscape, and decreased in periods 2 and 3 coinciding with an increase in the elephant population, but also to a decade of low dry season rainfall (Valeix et al. 2008). These patterns are coherent with the hypothesis that bottom-up mechanisms drive the patterns of densities we observe. Note that we did not explore other factors underlying the population trends of hyaena and its prey, such as the role of other large carnivores or climatic conditions, which was beyond the scope of this study. For example, the African lion, *Panthera leo*, which population has undergone large fluctuations, has the potential to influence herbivore populations and hyaena foraging behaviour (Grange et al. 2015, Périquet et al. 2015, Loveridge et al. 2016). Moreover, even though the hyaena population seems to decline 2-3 years after its prey population (Figure 3), we could not test for lags between their densities because periods were too short.

The main result of this study does not allow to conclude that a mechanism of hyper- or hypo-predation interfered with the population dynamics of predators and their prey on the long-term. Nonetheless some depicted patterns, although subtle, could be interpreted in the light of a role of scavenging. In period 1, both prey and hyaena population abundances increased, but this increase was stronger for hyaena (Table 1), which could be explained by hyaenas benefiting from both the increase in prey populations and the additional energy-rich resources provided by elephant culling. In period 2, growth rate confidence intervals are large and encompass 0 for both prey and hyaenas (Table 1), showing a general stabilization for the two populations. However, estimated growth rates suggest that prey abundance tended to decrease while hyaena abundance continued increasing in period 2, which is consistent with a scenario whereby the hyaena population, then abundant, compensated for the loss of the energy-rich resources provided by elephant carrion in period 1 by a renewed predation pressure in period 2. In period 3, hyaena population abundance did not decrease at a rate lower than its prey population, but maintains a higher density than at the start of period 1 despite a strong decrease of prey density, which could be explained by the presence of the additional food source provided by elephant carcasses. Finally, during period 4, both hyaena and prey densities increased, but the rate of increase is slightly lower for prey than hyaena, possibly indicating again a renewed predation of hyaena on prey populations. Differences between estimated growth rates of each guild are also potentially suggesting a change in the link between hyaena and prey densities caused by elephant carrion abundance : the differences are similar in periods of lower abundance of elephant carrion (about 0.06), but higher and lower in periods of high elephant carrion abundance. Testing these hypotheses would require further investigations in the future, by comparing hyaena diet and hunting behaviour in drought years (when elephant mortality will be high) and in years with good annual rainfall (when elephant mortality will be low).

Overall, these patterns are indicative of a numerical response of hyaena to the increased elephant carcass availability, through a rapid increase in density in period 1 and the maintenance of densities higher than at the start of our study period in period 3. If they are not hunted, mammalian predators’ density are generally associated with prey biomass (Fuller and Sievert 2001, Hayward et al. 2007) and this was verified for hyaena in Kruger National Park, South Africa (Ferreira and Funston 2016). Thus, a large increase of exploitable resources though inputs of additional food sources such as mega-carcasses should allow the survival and reproduction of more individuals; in other words, results in an increased carrying capacity (Oro et al. 2013). Contrary to our study, in Manitoba (Canada), the increased arctic fox (*Vulpes lagopus*) abundance after increased scavenging on carcasses of polar bear (*Usrus maritimus*) kills was accompanied by a signal in the population dynamics of its usual main prey, the collared lemming (*Dicrostonyx richardsoni*) (Roth 2003). The classical cyclic dynamics of lemming was delayed, probably due to an increased predation pressure arising from the higher predator abundance, which is akin to a phenomenon of hyper-predation (Courchamp et al. 2000, Moleón et al. 2014). However, arctic and African savannas ecosystems differ fundamentally. While arctic predator-prey communities are sometimes referred as the simplest in the world and present few predators that all rely largely upon one main prey, lemmings (*Dycrostony spp*., *Lemmus spp*.; Gilg et al. 2006, Reiter and Andersen 2011), African savannas present a high diversity of prey and predators in which classical prey and predator couplings are not observed (Sinclair et al. 2003). In arctic ecosystems, arctic fox only shifts to other prey when lemmings become very scarce (Gilg et al. 2006), which makes direct and observable repercussion of changes in their populations dynamics on that of lemmings more likely. In contrast, hyaena is known to be a generalist predator with a very plastic feeding behaviour, selecting food sources depending on their abundance (Kruuk 1972, Périquet et al. 2015). Consequently, if hyaena predation starts to reduce the abundance of one prey, it is likely that the predator shifts easily to another type of prey, allowing the other species to recover.

Furthermore, carrion can be considered as pulsed resources whose characteristics can affect the consumer dynamics (Yang et al. 2008). Interestingly, the spatio-temporal predictability of elephant carrion pulses differed between periods 1 and 3. During period 3, they were predictable and limited in time and space, as elephants die naturally near waterholes at the end of the dry season (Conybeare and Haynes 1984). From a hyaena perspective, a cursorial hunter, the end of the dry season is the time of the year when prey are the weakest and hence easier to catch (Cozzi et al. 2015). Consequently, mortality is likely to be compensatory, (meaning that individuals killed by hyaena were weak and likely to have died from other causes, such that population’s total mortality remained unchanged) and resource pulses at the end of the dry season may not matter that much. In contrast, during period 1, carcasses were highly aggregated (because entire elephant families were removed through culling) and unpredictable in space and time (Cumming 1981). They were more likely to influence hyaena fitness and hence demography with later consequences on prey demography. However, the unpredictability and short duration of carrion availability through these measures may explain the lack of strong and observable impacts on population dynamics on the long term as we observed here.

The frequency and intensity of pulses of carrion are likely to be altered under global changes and human activities. In large terrestrial mammals, peaks of herbivore carcasses are mostly associated with extreme weather periods, such as droughts or severe winters, which should become more frequent in the future (Intergovernmental Panel on Climate Change (IPCC) 2023). Moreover, anthropogenic carcasses become increasingly present (Moreno-Opo and Margalida 2019) through hunting activities (Mateo-Tomás et al. 2015), poaching (Underwood et al. 2013) or increasing number of domestic herbivores (Arrondo et al. 2019). Understanding how the characteristics of pulses of large herbivore carcasses may affect the coupling between large carnivores and their live prey populations is key if we want to predict the future of these populations.

## Acknowledgment

We are indebted to the Wildlife Environment Zimbabwe and thank warmly the Zimbabwe Parks and Wildlife Management Authority for their dedicated effort in monitoring wildlife in Zimbabwe in general and in Hwange National Park in particular and for providing this invaluable long-term dataset that spans over 50 years. We also thank Arnold Tshipa, Makalolo Camp and Wilderness for providing their rainfall data for this study. This manuscript benefited from comments from Eli Strauss and one anonymous reviewer, and a preprint version of this article has been peer-reviewed and recommended by PCIEcology.

## References

Abrams, P. A. 2000. The evolution of predator-prey interactions: Theory and evidence. - Annu. Rev. Ecol. Syst. 31: 79–105.

Andrén, H. et al. 2011. Modelling the combined effect of an obligate predator and a facultative predator on a common prey: lynx Lynx lynx and wolverine Gulo gulo predation on reindeer Rangifer tarandus. - Wildl. Biol. 17: 33–43.

Arrondo, E. et al. 2019. Rewilding traditional grazing areas affects scavenger assemblages and carcass consumption patterns. - Basic Appl. Ecol. 41: 56–66.

Bate, A. M. and Hilker, F. M. 2012. Rabbits protecting birds: Hypopredation and limitations of hyperpredation. - J. Theor. Biol. 297: 103–115.

Burnham, K. P. et al. 2011. AIC model selection and multimodel inference in behavioral ecology: some background, observations, and comparisons. - Behav. Ecol. Sociobiol. 65: 23–35.

Chamaillé-Jammes, S. et al. 2008. Resource variability, aggregation and direct density dependence in an open context: the local regulation of an African elephant population. - J. Anim. Ecol. 77: 135–144.

Child, G. 2004. Elephant culling in Zimbabwe. - ZimConservation Opin.

Conybeare, A. and Haynes, G. 1984. Observations on elephant mortality and bones in water holes. - Quat. Res. 22: 189–200.

Cortés-Avizanda, A. et al. 2009a. Carcasses increase the probability of predation of ground-nesting birds: A caveat regarding the conservation value of vulture restaurants. - Anim. Conserv. 12: 85–88.

Cortés-Avizanda, A. et al. 2009b. Effects of carrion resources on herbivore spatial distribution are mediated by facultative scavengers. - Basic Appl. Ecol. 10: 265–272.

Courchamp, F. et al. 2000. Rabbits killing birds: modelling the hyperpredation process. - J. Anim. Ecol. 69: 154–164.

Cozzi, G. et al. 2015. Effects of Trophy Hunting Leftovers on the Ranging Behaviour of Large Carnivores: A Case Study on Spotted Hyenas. - PLOS ONE 10: e0121471.

Cumming, D. 1981. The management of elephant and other large mammals in Zimbabwe. - In: Problems in management of locally abundant wild mammals. Academic Press, pp. 91–118.

DeVault, T. L. et al. 2003. Scavenging by vertebrates: behavioral, ecological, and evolutionary perspectives on an important energy transfer pathway in terrestrial ecosystems. - Oikos 102: 225–234.

Dudley, J. P. et al. 2001. Drought mortality of bush elephants in Hwange National Park, Zimbabwe. - Afr. J. Ecol. 39: 187–194.

Fallows, C. et al. 2013. White Sharks (Carcharodon carcharias) Scavenging on Whales and Its Potential Role in Further Shaping the Ecology of an Apex Predator - PLoS ONE 8: e60797.

Ferreira, S. M. and Funston, P. J. 2016. Population estimates of spotted hyaenas in the Kruger National Park, South Africa. - Afr. J. Wildl. Res. 46: 61–70.

Fuller, T. and Sievert, P. 2001. Carnivore demography and the consequences of changes in prey availability. - In: Carnivore Conservation. Cambridge University Press, pp. 163–178.

Gilg, O. et al. 2006. Functional and numerical responses of four lemming predators in high arctic Greenland. - Oikos 113: 193–216.

Gomo, G. et al. 2017. Scavenging on a pulsed resource: quality matters for corvids but density for mammals. - BMC Ecol. 17: 22.

Grange, S. et al. 2015. Demography of plains zebras (Equus quagga) under heavy predation. - Popul. Ecol. 57: 201–214.

Grosbois, V. et al. 2008. Assessing the impact of climate variation on survival in vertebrate populations. - Biol. Rev. 83: 357–399.

Guiden, P. W. et al. 2019. Predator-Prey Interactions in the Anthropocene: Reconciling Multiple Aspects of Novelty. - Trends Ecol. Evol. 34: 616–627.

Hayward, M. W. et al. 2007. Carrying capacity of large African predators: Predictions and tests. - Biol. Conserv. 139: 219–229.

Hebblewhite, M. et al. 2002. Elk population dynamics in areas with and without predation by recolonizing wolves in Banff National Park, Alberta. - Can. J. Zool. 80: 789–799.

Holmes, E. E. et al. 2012. MARSS: Multivariate Autoregressive State-space Models for Analyzing Time-series Data. - R J. 4: 11.

Holmes, E. E. et al. 2021. MARSS: Multivariate Autoregressive State-Space Modeling.

Intergovernmental Panel on Climate Change (IPCC) 2023. Climate Change 2021 – The Physical Science Basis: Working Group I Contribution to the Sixth Assessment Report of the Intergovernmental Panel on Climate Change. - Cambridge University Press.

Krebs, C. J. et al. 2001. What Drives the 10-year Cycle of Snowshoe Hares? - BioScience 51: 25.

Kruuk, H. 1972. The spotted hyena, a study of predation and social behavior. - Univ Chic. Press: 335.

Loveridge, A. J. et al. 2016. Conservation of large predator populations: Demographic and spatial responses of African lions to the intensity of trophy hunting. - Biol. Conserv. 204: 247–254.

Luna, Á. et al. 2021. Predation and Scavenging in the City: A Review of Spatio-Temporal Trends in Research. - Diversity 13: 46.

Mateo-Tomás, P. et al. 2015. From regional to global patterns in vertebrate scavenger communities subsidized by big game hunting. - Divers. Distrib. 21: 913–924.

Mellard, J. P. et al. 2021. Effect of scavenging on predation in a food web. - Ecol. Evol. 11: 6742–6765.

Moleón, M. and Sánchez-Zapata, J. A. 2015. The Living Dead: Time to Integrate Scavenging into Ecological Teaching. - BioScience 65: 1003–1010.

Moleón, M. et al. 2014. Inter-specific interactions linking predation and scavenging in terrestrial vertebrate assemblages. - Biol. Rev. Camb. Philos. Soc. 89: 1042–1054.

Moreno-Opo, R. and Margalida, A. 2019. Human-Mediated Carrion: Effects on Ecological Processes. - In: Olea, P. P. et al. (eds), Carrion Ecology and Management. Wildlife Research Monographs. Springer International Publishing, pp. 183–211.

Okabe, A. et al. 2009. Spatial Tessellations: Concepts and Applications of Voronoi Diagrams. - John Wiley & Sons.

Oro, D. et al. 2013. Ecological and evolutionary implications of food subsidies from humans. - Ecol. Lett. 16: 1501–1514.

Pereira, L. M. et al. 2014. Facultative predation and scavenging by mammalian carnivores: seasonal, regional and intra-guild comparisons: Predation vs. scavenging in carnivores. - Mammal Rev. 44: 44–55.

Périquet, S. et al. 2015. Spotted hyaenas switch their foraging strategy as a response to changes in intraguild interactions with lions. - J. Zool. 297: 245–254.

Périquet, S. et al. 2021. Dynamic interactions between apex predators reveal contrasting seasonal attraction patterns. - Oecologia 195: 51–63.

R Core Team 2021. R: A language and environment for statistical computing. - R Foundation for Statistical Computing, Vienna, Austria.

Reiter, M. E. and Andersen, D. E. 2011. Arctic Foxes, Lemmings, and Canada Goose Nest Survival at Cape Churchill, Manitoba. - Wilson J. Ornithol. 123: 266–276.

Rodewald, A. D. et al. 2011. Anthropogenic resource subsidies decouple predator—prey relationships. - Ecol. Appl. 21: 936–943.

Roth, J. 2003. Variability in marine resources affects arctic fox population dynamics. - J. Anim. Ecol. 72: 668–676.

Sinclair, A. R. E. et al. 2003. Patterns of predation in a diverse predator–prey system. - Nature 425: 288–290.

Slotow, R. et al. 2008. Lethal Management of Elephants. - In: Scholes, R. J. and Mennell, K. G. (eds), Elephant Management: A Scientific Assessment for South Africa. Wits University Press.

Underwood, F. M. et al. 2013. Dissecting the Illegal Ivory Trade: An Analysis of Ivory Seizures Data. - PLoS ONE 8: e76539.

Valeix, M. et al. 2008. Fluctuations in abundance of large herbivore populations: insights into the influence of dry season rainfall and elephant numbers from long-term data. - Anim. Conserv. 11: 391–400.

Wang, H. et al. 2009. The roles of predator maturation delay and functional response in determining the periodicity of predator-prey cycles. - Math. Biosci. 221: 1–10.

Wilson, E. E. and Wolkovich, E. M. 2011. Scavenging: how carnivores and carrion structure communities. - Trends Ecol. Evol. 26: 129–135.

Yang, L. H. et al. 2008. What can we learn from resource pulses? - Ecology 89: 621–634.

ZPWMA 1998. Hwange National Park Management Plan 1999-2004. Report, Harare, Zimbabwe.

