## Supplementary material for "Insights on the effect of mega-carcass abundance on the population dynamics of a facultative scavenger predator and its prey"

Appendix 1: Number of waterholes holding water at the end of the dry season, and hence monitored, in Hwange National Park (top) and in the study area (bottom) from 1972 to 2020. Dotted lines delimitate the different study periods: those with an elephant pictogram are the periods with a high abundance of elephant carcasses in the landscape (periods 1 and 3).

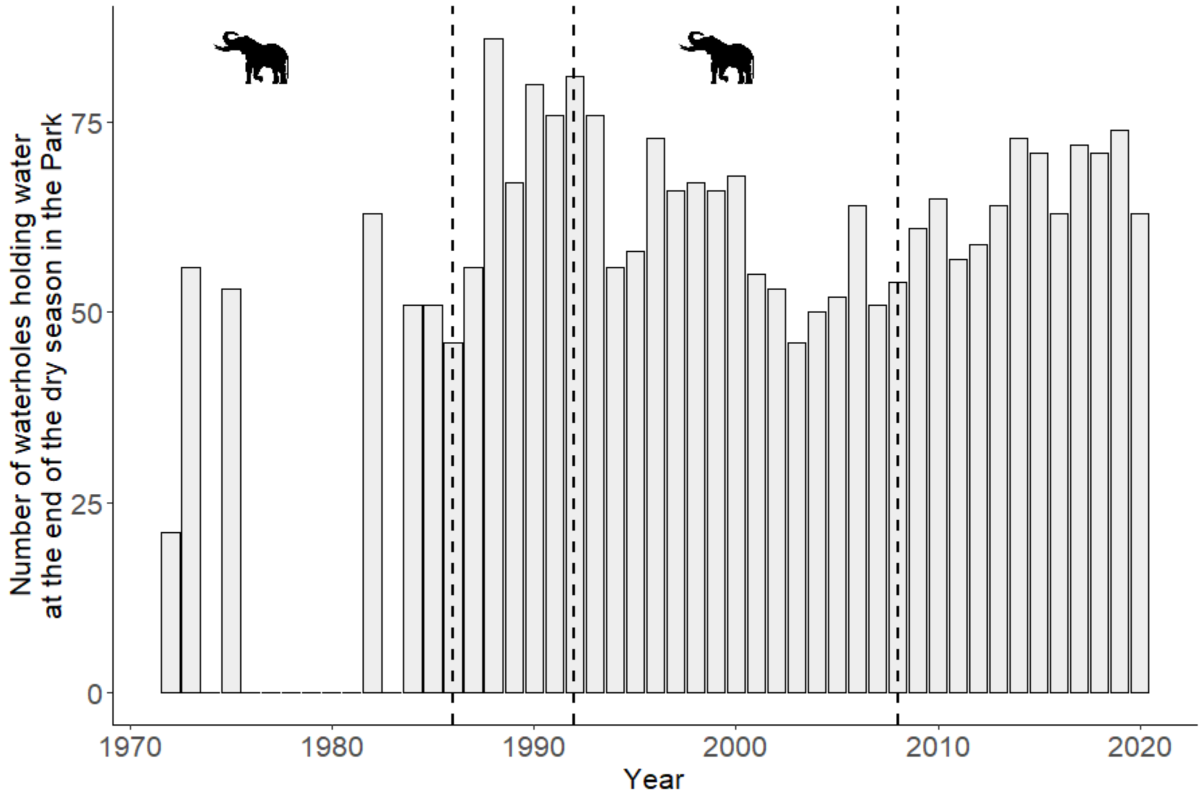


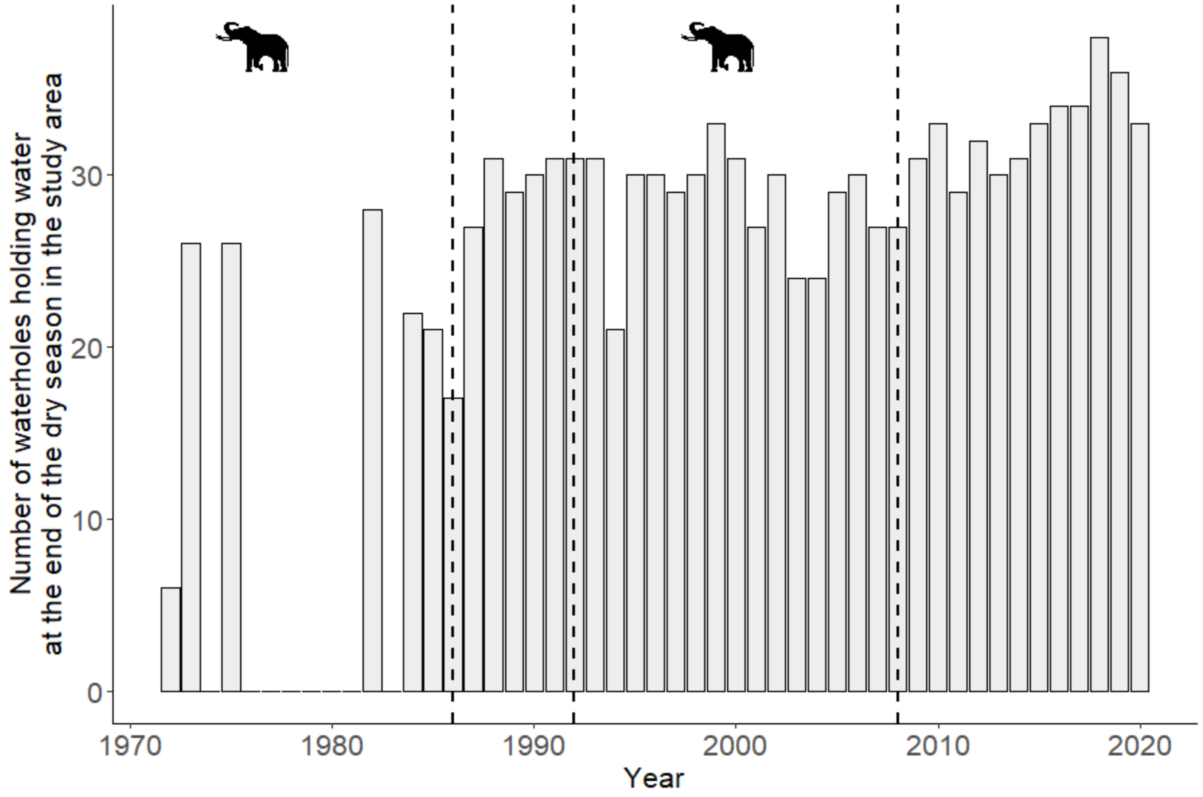

Appendix 2: Rainfall information.

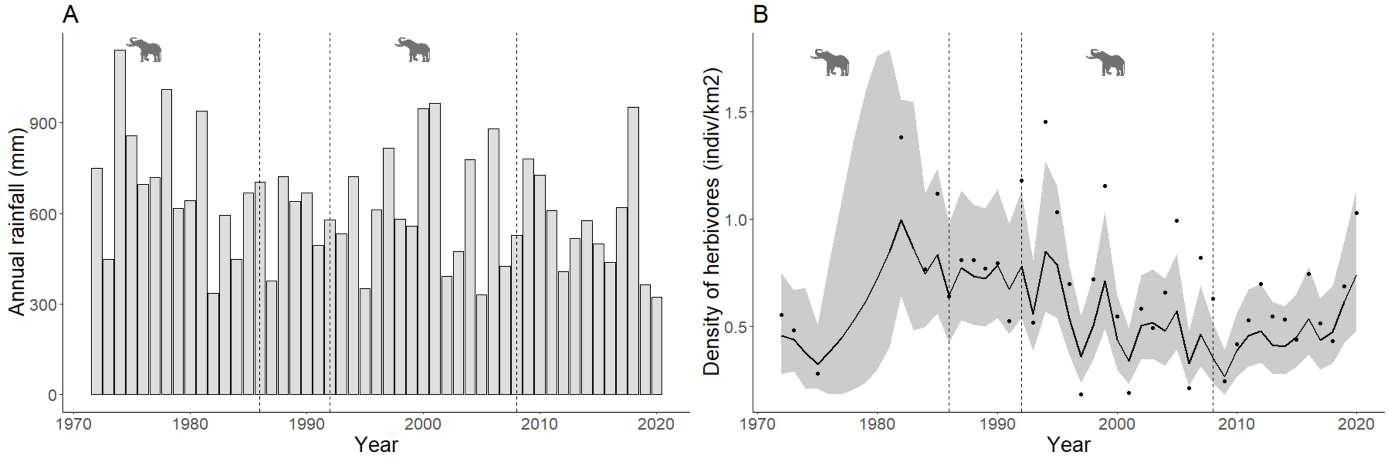


C

| Species | Model | AICc without rainfall | AICc with rainfall |
|---|---|---|---|
| Prey | Constant | 1892.00 | 1862.05 |
| | Time Varying | 1888.49 | 1890.89 |
| | Period | 1897.15 | 1850.05 |
| Hyaena | Constant | 3046.43 | 3039.18 |
| | Time Varying | 3086.64 | 3089.02 |
| | Period | 3037.43 | 3029.70 |

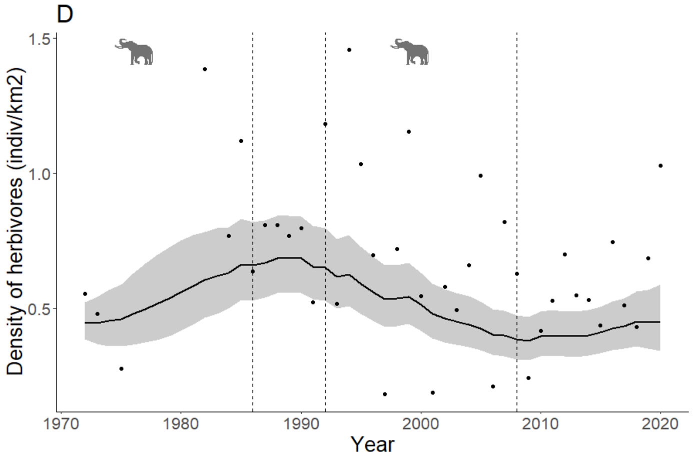


A) Annual rainfall in the study area. Data were collected from two meteorological stations: Main Camp until 2006 and Makalolo from 2007 to 2020. Annual rainfall for year y is calculated as the sum of the rainfall from October of year y-1 to September of year y. B) Density estimates of a constant model without annual rainfall as a covariate of the observation process. When annual rainfall is not considered, density estimates are overestimated in dry years, such as 2020, because animals need to visit waterholes more often to drink compared to wet years, such as 1974, for which density estimates are underestimated because animal need to visit waterholes less since natural ponds that allow them to drink are numerous and found throughout the park. C) AICc values of models with and without annual rainfall as a covariate of the observation process. AICc are lower for models with annual rainfall than models without, except for the time-varying model. This could be due to an over parametrization because time varying models present 54 parameters when annual rainfall is added and 53 without annual rainfall as a covariate. D) Density estimates of a constant model with annual rainfall as a covariate of the observation process. Density estimates are corrected: they are lower in dry years and higher in wet years, compared to the model without covariate in panel B. For all panels, the dotted vertical lines delimitate the different study periods: periods with an elephant pictogram are the ones with a high abundance of elephant carrion in the landscape (periods 1 and 3).
